# Intra- and inter-individual metabolic profiling highlights carnitine and lysophosphatidylcholine pathways as key molecular defects in type-2 diabetes

**DOI:** 10.1101/413203

**Authors:** Klev Diamanti, Marco Cavalli, Gang Pan, Maria J Pereira, Chanchal Kumar, Stanko Skrtic, Manfred Grabherr, Ulf Risérus, Jan W Eriksson, Jan Komorowski, Claes Wadelius

## Abstract

Type-2 diabetes (T2D) mellitus is a complex metabolic disease commonly caused by insulin resistance in several tissues. We performed a matched two-dimensional metabolic screening in tissue samples from a cohort of 43 multi-organ donors. The intra-individual analysis was assessed across five key-metabolic tissues (serum, adipose tissue, liver, pancreatic islets and muscle), and the inter-individual across three different groups reflecting T2D progression. We identified 92 metabolites differing significantly between non-diabetes and T2D subjects. Carnitines were significantly higher in liver, while lysophosphatidylcholines significantly lower in muscle and serum. An investigation of the progression to overt T2D showed that deoxycholic acid glycine conjugate was significantly higher in liver of pre-diabetes samples while additional increase in T2D was insignificant. A subset of lysophosphatidylcholines were significantly lower in the muscle of pre-diabetes subjects. Overall, the analysis of this unique dataset can increase the understanding of the metabolic interplay between organs in the development of T2D.

## INTRODUCTION

Type-2 diabetes (T2D) mellitus is a disease characterized by poor insulin sensitivity and failure of the pancreatic β-cells to secrete appropriate amounts of insulin^1,2^. Insulin resistance (IR) is associated with low physical activity, obesity, dyslipidemia and hypertension^3,4^. T2D is a major global health problem with a reported prevalence of 422 million in 2014 according to the World Health Organization (WHO)^5^. The insulin-related deficiencies in T2D lead to alterations in glucose, fatty acid uptake and metabolism, as well as lipid deposition in different tissues^6^.

The field of metabolomics has emerged through the development of high-throughput analytical chemistry techniques^2^. Metabolic profiling enables untargeted screening of a snapshot of the metabolic status for a given sample^7,8^. Liquid chromatography (LC) coupled with mass-spectrometry (MS) and gas chromatography (GC) coupled with MS are two prominent metabolic profiling technologies covering a wide spectrum of cellular low-weight compounds (<1500 Da)^7^. The untargeted screening is followed by computational “translation” of a subset of the identified compounds into known metabolites^9^.

IR in several tissues in combination with insufficient insulin secretion lead to systemic metabolic defects. Multiple studies have attempted to identify potential T2D biomarkers through metabolic profiling of blood, urine or saliva^1,6-8,10-12^. Risk factors identified in biofluids may partly reflect ongoing metabolic processes throughout the body, but are secondary to events occurring in other tissues^13^. Adipose tissue, liver, pancreatic islets and skeletal muscle are well-known to contribute to T2D development^14^, but others like brain and gut are also of importance.

In this study we performed a metabolic screening in a cohort of 43 organ donors. The study spanned across five key metabolic tissues (blood serum, intra-abdominal/visceral adipose tissue (VAT), liver, pancreatic islets and skeletal muscle) for T2D. The subjects were then in turn carefully matched for age, body-mass index (BMI), gender and per-tissue sample weight across the different phenotypes to accurately capture the metabolic fingerprint of T2D progression. To our knowledge, this is the first attempt to create an intra-individual map of metabolites across various metabolic-relevant tissues for the same cohort of individuals. In total we identified 286 unique metabolites across all tissues, 32% of which were significantly altered between non-diabetes and T2D. Amino acids (AAs) were elevated in VAT, liver, muscle and serum in T2D, and significantly associated to the percentage of glycosylated hemoglobin A1c (HbA1c). Bile acids were similarly higher, but only in liver. We showed that carnitines and lysophosphatidylcholines (LPCs) are the most significantly altered metabolites in muscle, liver and serum in T2D. Finally, we explored the progression to overt T2D and suggest potential biomarkers in the early development of T2D, i.e. from non-diabetes to pre-diabetes to T2D.

## RESULTS

### Metabolic profiling

We performed an untargeted metabolic profiling with GC-MS and LC-MS for 43 organ donors from healthy, pre-diabetes and T2D cohorts across five metabolic tissues: VAT, liver, pancreatic islets, skeletal muscle and blood serum (Figure 1). Samples were matched for BMI, age, gender and per-tissue sample weight (Table 1; Supplementary Table S1). The putative identity of the metabolites was determined computationally and resulted in 142 and 144 unique metabolites for GC-MS and LC-MS, respectively. The samples were classified in three distinct phenotypes related to the progression of T2D (controls, pre-diabetes and T2D). Partial least squares – discriminant analysis (PLS-DA) models showed a higher similarity of pre-diabetes to controls when compared to T2D, hence controls and pre-diabetes subjects were merged into one non-diabetes group and compared to T2D (Table 1; Supplementary Figures S1A-B, S2A-B). In total 92 unique metabolites varied significantly in at least one tissue between non-diabetes and T2D samples (Figure 2).

**Table 1:** Baseline characteristics of the 43 subjects in the cohort. The mean value and the standard deviation are shown for age, BMI, HbA1c and GSIS. Gender is shown as the proportion of females (F) and males (M). Age is expressed in years. BMI is in kg/m^2^. HbA1c is in % mmol/mol. GSIS is mmol in liters of glucose. Non-diabetes is the merged group of controls and pre-diabetes. P-value 3C is the p-value calculated from a Kruskal-Wallis test on controls, pre-diabetes and T2D. P-value 2C is the p-value calculated from a Wilcoxon-Mann Whitney test on non-diabetes and T2D.

| *Parameter* | *Controls* | *Pre-diabetes* | *Non-Diabetes* | *T2D* | *P-value 3C* | *P-value 2C* |
| --- | --- | --- | --- | --- | --- | --- |
| *Gender* | 7 F / 10 M | 4 F / 9 M | 11 F / 19 M | 4 F / 9 M | - | - |
| *Age* | 59 ± 10 | 64 ± 8 | 61 ± 9 | 65 ± 7 | 0.2124 | 0.3013 |
| *BMI* | 26.5 ± 3.8 | 27.6 ± 5.6 | 27.0 ± 4.6 | 27.9 ± 5.6 | 0.8417 | 1 |
| *HbA1c* | 5.4 ± 0.2 | 5.9 ± 0.1 | 5.6 ± 0.3 | 7.3 ± 1.4 | 1.1×10<sup>-8</sup> | 6.9×10<sup>-7</sup> |
| *GSIS* | 11.8 ± 5.6 | 15.1 ± 30.8 | 13.2 ± 20.3 | 5.0 ± 2.5 | 1.3×10<sup>-3</sup> | 2.6×10<sup>-3</sup> |

**Figure 1:**
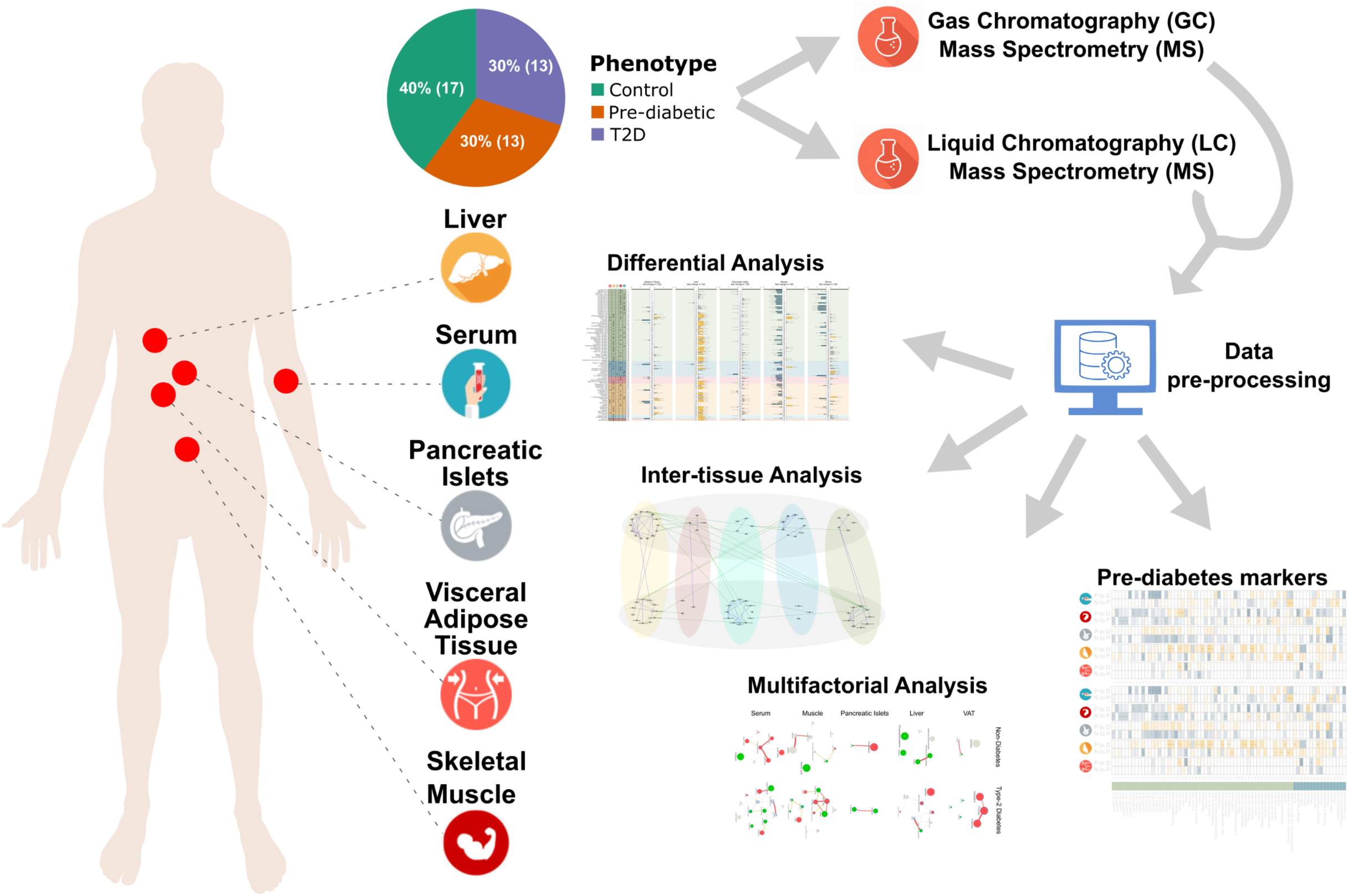
Schematic overview of the study summarizing sample collection, number of subjects, metabolic profiling, data processing and computational analysis.

**Figure 2:**
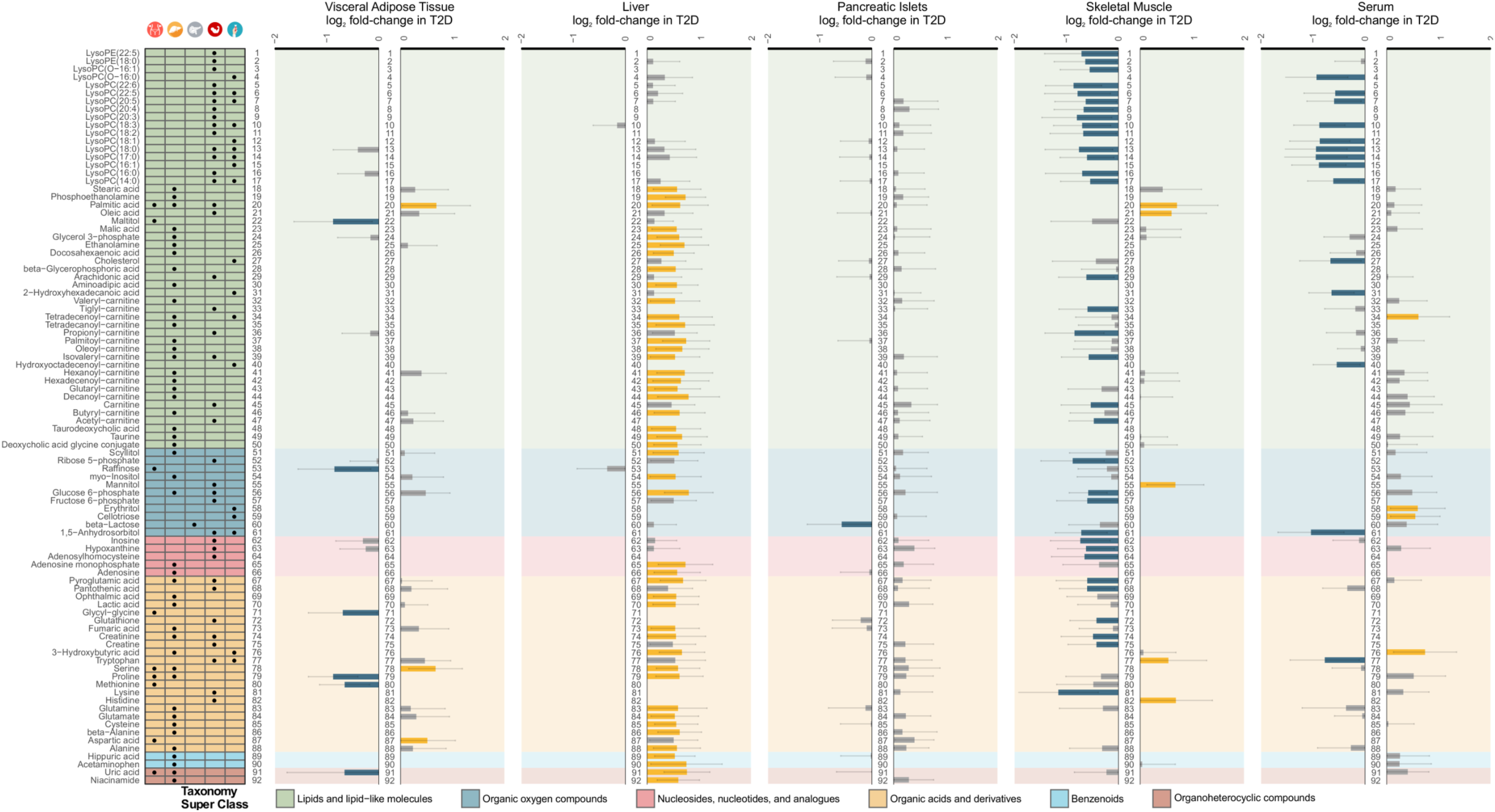
Overview of the differential and the fold-change analysis of the computationally annotated compounds from GC-MS and LC-MS for non-diabetes versus T2D subjects. Rows of the table on the left-hand-side contain the 92 metabolites that were significant in at least one tissue. Table columns represent each of the five tissues (VAT, liver, pancreatic islets, skeletal muscle and serum). A black dot implies statistical significance in the corresponding tissue (p-value<0.1). The color-coding in the table originates from a curated classification of the Human Metabolome Database (HMDB) super-class taxonomy and the labels are explained in the legend^51^. The five barplots represent the log_2_ fold-change in T2D of the μ=0 and σ=1 scaled log_2_ compound intensities are in the same order as the tissues in the table. Error bars represent the 90% confidence intervals. Yellow bars imply statistical significance and increase, blue bars statistical significance and decrease, while grey bars did not cross the statistical significance threshold. Numbering is to assist following the variation of metabolite across tissues.

Intensities of internal standards (IS), deviation of experimental retention time and mass from their theoretical values and principal component analyses (PCAs) for the MS running order of samples were used to assess the quality of the profiling (Supplementary Note – Overview of metabolic profiling). IS varied mainly in LC-MS for VAT (Supplementary Figures S3 and S4). However, the relative standard deviation of the intensities in LC-MS was below 20% for all IS that ionize well in the respective mode. PCAs showed no evidence for effect of the running order (Supplementary Figures S5 and S6). Divergence of experimental retention time and mass for IS from the expected ones were within accepted limits (Supplementary Figures S7 and S8).

### Alterations of metabolites across tissues in T2D

After imputing and transforming the intensities of the metabolites we investigated the partitioning of the samples per tissue (Materials and methods – Data transformation and normalization). Large within-group variation led PCAs towards ambiguous separation (Supplementary Figure S9A), yet orthogonal PLS-DA ((O)PLS-DA) improved the partitioning of the phenotypic groups (Supplementary Figure S10A). Covariates, excluding HbA1c and glucose-stimulated insulin secretion (GSIS), were suggested (cf. Methods - Data transformation and normalization; Supplementary Table S2; Supplementary Figure S11) and proven (Supplementary Table S3; Supplementary Figures S9B and S10B) to maintain the separation seen in the scaled and log_2_-transformed raw intensities (Supplementary Figures S9A and S10A). Correcting for BMI, age, gender and sample weight did not yield extensive alterations in the set of significantly varying metabolites across tissues (Supplementary Figure S12B).

Next, we sought to explore the statistical significance and the direction of the variation of the computationally annotated metabolite intensities between non-diabetes and T2D (Figure 2). More than half of the total number of significant metabolites belonged to the taxon of lipids and lipid-like molecules, which mainly consisted of LPCs, carnitines and free fatty acids (FFAs). Specifically, LPCs were overrepresented in muscle and serum (p-value<10^-132^), while carnitines were abundant in liver (p-value<10^-132^) (Supplementary Table S4; Supplementary Figure S13). We further observed that LPCs and carnitines in serum followed similar patterns with muscle and liver, respectively (Figure 2). Various FFAs were noticed as significant among liver, muscle and serum (Figure 2).

Other important groups of metabolites were those of AAs and nucleosides, nucleotides and analogues. We additionally observed that glucose 6-phosphate levels were significantly higher in VAT and liver, while in muscle they were significantly lower, similarly to fructose 6-phosphate (Figure 2). These metabolites, together with ribose 6-phosphate that is also significantly lower in muscle of T2D, are part of the pentose phosphate pathway that has been extensively linked to T2D^15^.

### Regulation of carnitine and choline pathways

Carnitine represents a crucial *trait d’union* between carbohydrate and fatty acid (FA) metabolism. Carnitine facilitates the transport of FAs into the mitochondria for β-oxidation, but it is also involved in glucose metabolism regulating the pool of acetyl-CoA in the cytoplasm.

In our samples, we observed a significant increase in the pool of short- and medium-chain carnitines in liver in T2D and a similar trend in pancreatic islets and serum. Instead, a trend towards reduction of carnitines was observed in muscle and in particular very short-chain carnitines were significantly reduced in T2D (Figures 2 and 3).

**Figure 3:**
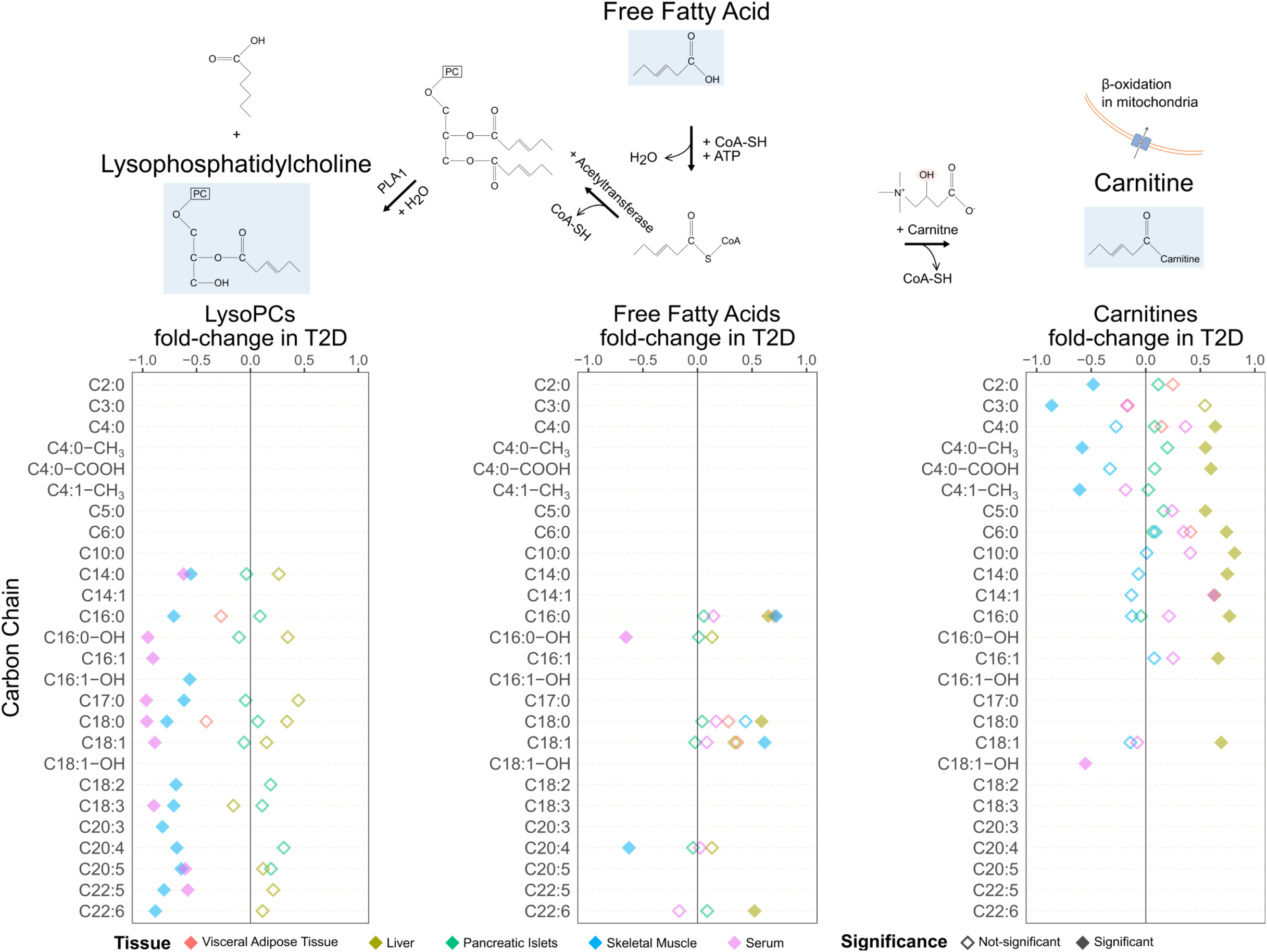
Schematic overview of the formation of LPCs and carnitines from FFAs. The top graph shows an example of the pathway of LPCs and carnitines formation as conjugates of FFAs (e.g. C6) with various other compounds via different enzymes. The three bottom plots describe the log_2_ fold-change and significance of LPCs, FFAs and carnitines. Rows represent carbon-chain length of the metabolites (Supplementary Table S7). Color-coding of the rhombuses represent tissues while the filling of the rhombus implies statistical significance.

The complexity of the molecular mechanisms behind the association of carnitines to T2D was highlighted using a systems biology approach by Bene et al^16^. They showed that individuals with T1D or metabolic syndrome share similar levels of medium- and long-chain acyl-carnitines while individuals with T2D or metabolic syndrome have similar levels of short-chain acyl-carnitine.

We also found a generalized reduction in LPC levels in both serum and muscle including LPC with a surprisingly wide variety of saturation and desaturation; C14:0, C16:0, C16:1, C17:0, C18:0, C18:1, C18:2, C18:3, C20:4, C20:5, C22:5 and C22:6 (Figure 3). The extent of decrease in LPCs is larger in muscle than in plasma (Figure 2).

### Increased drive of bile acid pathway in T2D

We observed a statistically significant increase in the levels of deoxycholic acid conjugate with glycine or taurine in liver in T2D individuals, whereas the increase of chenodeoxycholic acid conjugates was not reaching statistical significance (Figure 2). Glycodeoxycholate was also higher in liver, muscle and serum, but to a significant level only in liver. Taurine levels were also significantly higher in liver of T2D individuals with a non-statistically significant trend also in pancreatic islets, muscle and serum (Supplementary Figure S14). Taurine, which has been reported to associate to serum glucose levels in animal models^17^, is synthesized in the pancreas from cysteine, an AA we detected as significantly elevated in T2D liver samples (Figure 2; Supplementary Figure S15). The pathways of primary bile acids biosynthesis and taurine-hypotaurine metabolism were significantly enriched in liver (p-value<2.18×10^-2^) (Supplementary Table S5; Supplementary Figure S16).

### AAs vary across tissues in T2D

High levels of circulating branched-chain AAs (BCAAs) leucine, isoleucine, valine and the aromatic AAs (AAAs) tryptophan, phenylalanine and tyrosine, have been repeatedly reported in the literature as associated to obesity, impaired fasting glucose and T2D^18^. Although the differences in BCAA and AAA did not cross the statistical significance threshold levels between control and T2D individuals, we observed a higher level of valine and isoleucine in serum in T2D (Supplementary Figure S17). The same trend was observed in VAT. In liver and pancreatic islets the levels of valine, isoleucine and leucine were higher in T2D, while in muscle BCAAs were depleted in T2D (Supplementary Figure S17). Tyrosine levels were elevated in VAT, muscle and serum and phenylalanine was higher in T2D liver, pancreatic islets and VAT samples (Supplementary Figure S17).

Adams and colleagues utilized plasma metabolite patterns to target muscle-specific metabolites in African-American females with significantly reduced whole-body lipid oxidation^6^. They found that higher levels of leucine and valine were strongly correlated with acetyl-carnitine, which we found significantly lower in muscle in T2D (Figures 2 and 3). Here we observed leucine, isoleucine and valine to be strongly correlated to acetyl-carnitine in muscle (Spearman correlation r>0.65, p-value<0.0102), which was also correlated to propionyl-carnitine (Spearman correlation r=0.64, p-value<0.0161), a marker of valine’s catabolic product propionyl-CoA^6,10^.

Linear regression models showed that HbA1c was associated to various BCAAs and AAAs for VAT, liver, muscle and serum. Specifically, we observed that tyrosine, phenylalanine, leucine and isoleucine in VAT were significantly associated with HbA1c (Figure 4A). Tyrosine and leucine shared the same significant pattern in VAT and serum (Figure 4A; Supplementary Figure S17). Interestingly tryptophan was significant for muscle and serum in T2D (Figure 2), and significantly associated with HbA1c in muscle, liver and serum, sharing an increasing pattern in liver and muscle, while decreasing in serum (Figures 2 and 4A; Supplementary Figure S15). Elevated levels of tryptophan in serum is associated to T2D, making it a potential risk predictor for the pathogenesis of T2D^19^. There is increasing evidence pointing towards links between inflammatory processes and tryptophan-related metabolites, e.g. kynurenine that was higher in VAT, liver and muscle of T2D, but did not cross significance threshold (Supplementary Figure S15)^20^.

**Figure 4:**
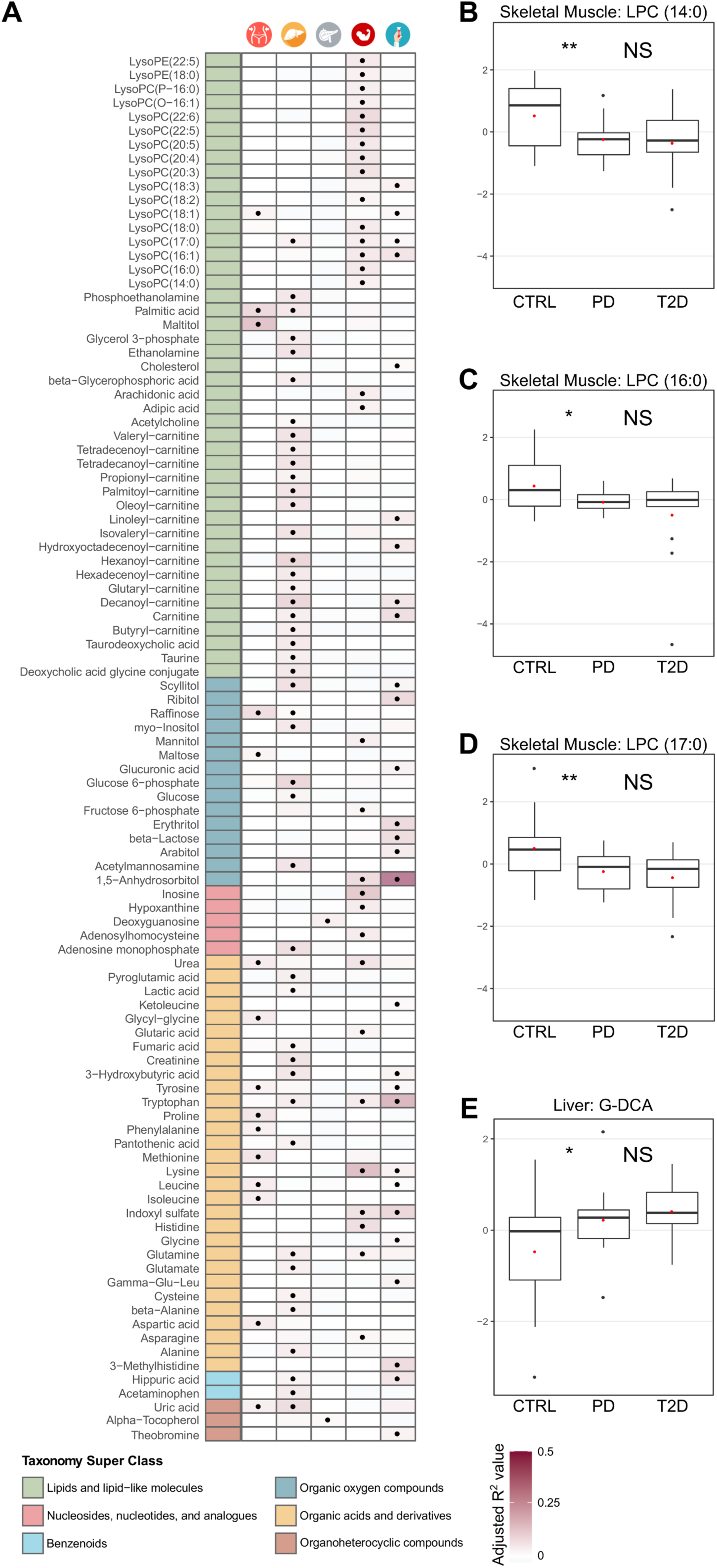
A) Overview of the metabolites that are significantly linearly associated with HbA1c in at least one tissue. The first column is color-coded according to a curated classification of the HMDB super-class taxonomy and the labels are explained in the legend. The following five columns represent each tissue as noted on the top of the heatmap. A black dot implies statistically significant association of the corresponding metabolite to HbA1c (p-value<0.1). The color intensity in the cell background shows the level of the adjusted R^2^ value from the linear regression model. **B–E)** Levels of selected metabolites for pair-wise comparisons among controls (CTRL), pre-diabetes (PD) and T2D subjects. Statistical significance is shown for each pair-wise comparison (p-value>0.1 NS; 0.1≥p-value>0.05 *; 0.05≥p-value>0.01 **; 0.01≥p-value>0.001 ***; 0.001≥p-value ****) (cf. Methods – Statistical analysis). The red dot signifies the mean value of the group. **B)** LPC (14:0) in skeletal muscle; **C)** LPC (16:0) in skeletal muscle; **D)** LPC (17:0) in skeletal muscle; **E)** Deoxycholic acid glycine conjugate (G-DCA) in liver.

We also took advantage of the recent development of an R tool MoDentify^21^ to scan for metabolites correlated to the phenotype as single entities or as members of functional modules^22^. This analysis on the full set of AAs revealed phenotype-driven functional modules, where single metabolites that are not directly correlated to T2D form combinatorial modules which are significantly associated to the phenotype. A large number of BCAAs and AAAs, together with other AAs, that are significant either individually or as part of modules were shown to be associated with T2D. They include, among others, leucine, isoleucine, tyrosine and glycine (Supplementary Figure S18).

Various other AAs have been associated to pre-diabetes or T2D with less consistency^23^. Particularly higher levels of alanine, serine, proline, glutamine, the glutamine/glutamate ratio, glutamate, histidine and lysine. Here we observed significantly higher levels of alanine, glutamine, glutamate, proline and serine in liver; significantly lower levels of proline and higher levels of serine in VAT; and significantly higher levels of histidine in muscle in T2D (Figure 2; Supplementary Figure S15). Higher levels of glutamate have been linked to diabetes due to its function as a substitute energy source to glycolysis or β-oxidation^2^.

Significantly lower levels of methionine and lysine were observed in T2D VAT and muscle, respectively (Figure 2; Supplementary Figure S15). Methionine and lysine are precursors involved in the endogenous synthesis of L-Carnitine which was also significantly lower in T2D muscle tissues (Figure 2).

### FA-related pathways are enriched in T2D across tissues

Metabolite classification analysis revealed that VAT, liver, muscle and serum are enriched for AAs and various types of lipids. Specifically, VAT and serum were enriched for AAs (p-value<1.15×10^-3^). Liver and muscle were enriched for AAs and derivatives (p-value<1.13×10^-7^), and acyl carnitines (p-value<2.3×10^-7^). Muscle was highly enriched for LPCs, similar to serum (p-value<10^-132^) (Supplementary Table S4; Supplementary Figure S13). A pathway enrichment analysis showed various pathways related to lipids and lipid-like molecules, such as glycerophospholipid metabolism in liver (p-value<4.63×10^-4^), biosynthesis of unsaturated FAs in muscle (p-value<1.1×10^-3^) and carnitine synthesis in serum (p-value<1.59×10^-2^) (Supplementary Table S5; Supplementary Figure S16). Overall, liver was the tissue with the largest number of significant metabolic pathways, followed by muscle, VAT and serum (Supplementary Figure S19).

Liver, muscle and serum showed a strong enrichment in biological roles that are strongly connected to functionalities of lipids and lipid-like molecules in cells, such as lipid catabolism, fatty acid transport, energy production (p-value<9.48×10^-5^). Muscle metabolites were enriched in unsaturated fatty acids (p-value<1.99×10^-2^), and together with serum in membrane components and energy sources (p-value<8.09×10^-2^) (Supplementary Table S6; Supplementary Figure S20). VAT, liver and muscle also showed enrichment for essential AAs (p-value<4.03×10^-10^), and for various other AA-related biological roles (p-value<1.08×10^-3^) (Supplementary Table S6; Supplementary Figure S20).

We used MoDentify to confirm that similar metabolites and classes of metabolites were correlated^21^ (Supplementary Figures S21 and S22). Interestingly, we also observed a higher correlation of metabolites within tissues, rather than among tissues (Supplementary Figure S23). The most enriched correlation was that of metabolites belonging to the class of lipid and lipid-like molecules (Supplementary Figures S23 and S24).

### Rule-based classification suggests combinations of metabolites that define type-2 diabetes patients

We performed a multivariate statistical analysis to verify metabolites identified by the earlier univariate analysis and to discover new metabolites that were non-linearly associated with T2D in various tissues. Indeed, a subset of LPCs was also marked as significant by the multivariate statistical approach in muscle and serum. In agreement with the previous analysis, proline was found significant in liver, while lysine and histidine in muscle. Proline was also marked as significant for muscle, even though the univariate approach did not mark it as such. Threonine and asparagine were found as significant in discerning between non-diabetes and T2D in VAT (Supplementary Figure S25).

We next used the set of “all-relevant” selected metabolites from above as input to the ROSETTA machine-learning approach that produces transparent classification models to uncover combinations of variables that discriminate between phenotypes^24,25^. The computational models for liver, muscle and serum were of high quality (accuracy>80%), while those for VAT and pancreatic islets performed adequately well (Supplementary Table S8) given the complexity of the data and the sample size. Rough-set classification models consist of IF-THEN rules, which in addition to being legible, can be visualized into networks (Supplementary Table S9). Such networks show combinations of metabolites and their levels that differ between T2D and non-diabetes. In muscle combinations of high levels of palmitic acid and histidine, together with low levels of lysine and LPC(22:6) predict T2D, while average levels of LPC(22:6), mannitol and lysine predict non-diabetes (Supplementary Figure S26). In liver various levels of adenosine monophosphate in combination with other AAs such as proline and alanine play a central role in deciding for T2D or non-diabetes (Supplementary Figure S26).

### Progression of T2D through pre-diabetes

In order to identify potential early markers of T2D we applied two different approaches. We used the group of pre-diabetes to monitor the development of metabolites through pair-wise comparisons of the significance between controls and pre-diabetes, as well as between pre-diabetes and T2D. We also expressed HbA1c, a marker of pre-diabetes, as a function of various metabolites via linear regression models (cf. Materials and methods).

We first explored the separation among the three phenotypes through PCA and (O)PLS-DA, and observed that the separation was not satisfactory (Supplementary Figures S1 and S2). This occurred mainly due to the limitations in statistical power and the large variation of the metabolite levels. The analysis of T2D and the group of pre-diabetes revealed that 45% of the significantly altered metabolites belonged to the class of lipids and lipid-like molecules (Supplementary Figure S27A). A large proportion of these metabolites were significant when comparing pre-diabetes to T2D, but not when comparing controls and pre-diabetes. Liver showed a strong enrichment in significantly increased carnitines, FFAs and various AAs in T2D, which were also associated with HbA1c (Figure 4A; Supplementary Figure S27A). Compounds belonging to the class of nucleosides, nucleotides and analogues in muscle and six different LPCs in serum were significantly lower in T2D (Supplementary Figure S27A). We also observed an extended association of various AAs, including several BCAAs and AAAs, with HbA1c (Figure 4A).

In contrast to serum, various LPCs and FFAs in muscle showed a significant decrease in pre-diabetes when compared to controls, while the same compounds did not further alter or decrease in T2D (Figure 4B-D; Supplementary Figure S27A). Such a large collection of events of deregulation in pre-diabetes in combination with the large proportion of LPCs associated with HbA1c suggested that specific LPCs in muscle might be early predictors for T2D (Figure 4A-D). In serum and liver, it is also worth to observe that the levels of the bile acids pool increased between control and pre-diabetes while maintaining an elevated level in T2D subjects (Figure 4E; Supplementary Figure S27A). Changes in taurodeoxycholic acid, taurine and deoxycholic acid glycine conjugate were also significantly associated to the HbA1c values in the liver (Figure 4A). This provides additional evidence that imbalance of the cholesterol metabolism could represent an early marker of T2D.

## DISCUSSION

We performed an extensive metabolomics analysis in a unique collection of five tissues of key importance for T2D. To our knowledge, this is the first attempt to investigate such a wide range of metabolic profiles across tissues in T2D. In total 32% of all the detected metabolites were significant in at least one tissue when comparing non-diabetes to T2D. Generally, the levels of metabolites in liver and muscle, the two major insulin sensitive tissues, demonstrated divergent patterns, with metabolites in liver mainly increasing and the ones in muscle decreasing, potentially reflecting impaired tissue-specific IR and/or metabolic control (Figure 2). We also observed a total increase of BCAAs and their associations with HbA1c, previously connected with IR in various tissues^18^ (Figure 4A; Supplementary Figures S15 and S17). Reduced alanine levels in muscle and higher in liver in T2D, suggested an overall increase in the glucose-alanine cycle (Figure 2; Supplementary Figure S15).

Various AAs, including BCAAs and AAAs, were significantly associated with HbA1c in VAT which clinically, is more interesting and relevant to T2D than subcutaneous adipose tissue (SAT). The VAT depot is overall more closely linked to IR and T2D phenotype, with potential causal effects on disturbing hepatic glucose metabolism in prediabetes^26^. Petrus et al. have earlier shown that adipose tissue depots differ in lipid and fatty acid composition, which may be due to distinct enzymatic activity, lipolytic function or gene expression^27^. Hence, our findings in VAT may not apply to other adipose depots such as SAT.

The largest class of metabolites that was altered in T2D samples was that of lipid and lipid-like molecules, with LPCs and carnitines highly enriched, and sharing patterns seen in serum with muscle and liver, respectively (Figures 2 and 3). Plasma LPC levels are determined by lecithin-cholesterol acyltransferase (LCAT) activity, hepatic secretion and phospholipase A2 synthesis from PC^28^. Reduction of LPCs in plasma has been reported to be tightly associated with obesity^28,29^ or IR^1,30-32^. The observed systematic decrease in LPCs in both muscle and plasma when comparing the T2D and control subjects may be a strong indicator of diabetes progression (Figures 2 and 3). Interestingly, here we detected LPCs to significantly decrease in muscle of pre-diabetes and to further decrease in T2D, even though insignificantly (Figure 4B-D; Supplementary Figure S27A). Bruce and colleagues examined the impact of high-fat diet in LPCs of mice and detected similarly lower levels in the first week, indicating potential acute alterations^29^.

The endogenous carnitine pool is for the major part located in skeletal muscle. Carnitines and acyl-carnitines are also present in the gastrointestinal tract, the liver and the kidneys. Metabolic changes can affect the carnitine pools in the various tissues, but the homeostasis can vary between tissues so that for example, changes in carnitine content in liver rapidly appear in plasma, whereas changes in skeletal muscle content may not be as readily detected^33^. Most of the metabolic studies of the correlation between carnitines, IR and T2D so far relied on analyses of urine, serum and plasma. Here we extended the analysis of the carnitine pool in T2D in other relevant tissues such as liver, muscle, VAT and pancreatic islets (Figures 2, 3 and 4A).

Alterations in the levels of long and medium chain acyl-carnitines, including octadecenoyl-carnitine (C18:1), tetradecenoyl-carnitine (C14:1), tetradecadienoyl-carnitine (C14:2) and short chain acyl-carnitines, such as malonyl-carnitine/hydroxybutyryl-carnitine (C3DC+C4OH), have been reported both in T2D and pre-diabetes states in serum^3,10,34,35^. Here we observed a general increase in the short and medium-chain carnitines in liver in T2D, and a similar pattern in serum and pancreatic islets (Figures 2, 3 and 4A; Supplementary Figure S27A). Accumulation of long-chain acyl-carnitines and acyl-carnitine-species in T2D plasma might reflect an increased, yet incomplete, mitochondrial β-oxidation with even-chain acyl-carnitines, up to 20 carbons, resulting from the initial rounds of oxidation and odd-chain species associated to AA catabolism. The deregulation of β-oxidation seems to be unrelated to the activity of the carnitine acyltransferase I (CPT1) enzyme^36^. Additionally, 3-hydroxybutyric acid, a member of the ketone bodies, whose increase is known to be associated with enhanced β-oxidation, was higher in muscle and significantly higher in liver and serum (Figure 2)^37,38^.

Bile acids have been associated with the regulation of cholesterol catabolism, lipid-absorption, homeostasis of triglycerides and glucose. They can also act as hormones regulating various metabolic processes. T2D has been linked to variation of the bile acids pool composition, in particular the ratio of the pool of cholic and deoxycholic acid in the liver to chenodeoxycholic acid in the intestine is reported to be a metabolic signature of impaired cholesterol catabolism in T2D. Evidence in animal models and human points to a higher level of cholic acid synthesis in T2D, which in turn increases the level of deoxycholic acid^4,39,40^. Here we observed higher levels of bile acids in liver, muscle and serum of T2D (Figure 2; Supplementary Figure S14). A closer look at the levels of bile acids revealed strong correlations to HbA1c and increased levels in liver and serum in pre-diabetes and T2D (Figures 4A and 4E; Supplementary Figure S27A).

Carbohydrate metabolites such as glucose- and fructose-6-phosphate are increased in VAT, liver and serum, but reduced in muscle in T2D. This might reflect reduced metabolism in muscle due to IR and increased supply to other organs (Figure 2). Decanoyl-carnitine and deoxycholic acid glycine conjugate show the same significantly increased pattern in serum and liver of pre-diabetes (Supplementary Figure S27A). These serum markers might reflect their corresponding levels in liver as potential early markers for IR.

Despite the novel findings on metabolite changes across several tissues, this study has clear limitations. Due to ethical restrictions, we did not have access to all confounding factors. Hence, there are likely effects due to the acute disease conditions as well as medications and other interventions at the intensive care unit, including metabolic effects of stress. The results need to be confirmed in other settings, e.g. in cohorts of deceased donors with a larger set of potentially confounding factors. There was also lack of statistical power due to the limited collection of samples and, moreover, causality could not be inferred. Instead we explored sets of potential biomarkers that differed between pre-diabetes or T2D. Increasing the number of samples would partly overcome this limitation, and allow future investigations of the dysregulated molecular mechanisms. Further limitations are due to the complex extraction of metabolites from VAT, the relative compound quantification from MS and the uncertainty of the computational pairing of peaks to compounds. However, these limitations are expected to be addressed together with the constant advancements in the field of MS and standardization of complex sample preparation protocols.

## MATERIALS AND METHODS

### Ethics statement

The consent to use the organs of donors for scientific research was obtained from an online database that is documented to be in full accordance with the regional standard practices and the Swedish law or verbally by the physician in charge from the closest relatives of the deceased person. The tissue samples were acquired and stored at the Excellence of Diabetes Research in Sweden (EXODIAB) biobank. EXODIAB provided the samples that we further stored and analyzed in agreement with the Uppsala Regional Ethics Committee approval (Dnr: 2014/391).

### Sample collection and processing

Samples from three different phenotypic groups (healthy controls, pre-diabetes and T2D) were acquired from frozen portions of five metabolically-relevant human tissues (intra-abdominal/visceral adipose tissue (VAT), liver, pancreatic islets, abdominal skeletal muscle and blood serum) from the EXODIAB biobank. The distribution of the individuals and the tissues over the phenotypes is shown in Figure 1. Samples were matched for BMI, age, gender and per-tissue sample weight (Table 1; Supplementary Table S1). T2D patients were identified based on their medical records at the corresponding medical facilities according to the WHO guidelines^5^. Pre-diabetes patients were defined by the percentage of HbA1c level (5.7 % mmol/mol ≤ HbA1c ≤ 6.5 % mmol/mol) in blood (Table 1). Samples with HbA1c<5.7 % mmol/mol were classified as controls (Table 1). Controls and pre-diabetes were merged into one group (non-diabetes) into a primary analysis, and their metabolic profiling was compared to the one of T2D. Merging was decided based on PLS-DA models that showed a higher similarity of pre-diabetes to controls when compared to T2D, and for gain of statistical power (Supplementary Figures S1A-B). We also measured the GSIS in pancreatic islets that proved to be correlated to be impaired in T2D patients^41^ (Table 1; Supplementary Table S1; Supplementary Figure S11). In secondary analyses, we also examined all three groups separately.

### Metabolic profiling

Metabolic profiling with GC-MS and LC-MS was performed at the Swedish Metabolomics Center in Umeå, Sweden. Detailed information about sample preparation, mass spectrometry and targeted data processing is available at (Supplementary Note - Metabolic Profiling). The remaining supernatants of each tissue type were pooled and used to create tissue-specific quality control (QC) samples. Tandem mass spectrometry analysis (LC-MS only) was run on the QC samples for identification purposes. The samples were analyzed in per-tissue batches according to a randomized run-order on both GC-MS and LC-MS.

### Data transformation and normalization

log_2_ transformation was applied to the whole dataset after missing values were replaced by 1.00001 to avoid negative values. The transformation assisted in achieving a better approximation of a normal distribution and avoiding bias by outliers^7^. Quantifications from GC-MS and LC-MS were merged into one dataset and metabolite intensities were scaled to μ=0 and σ=1^42^.

ISs from MS were also included to the set of covariates, while HbA1c and GSIS were excluded due to their correlation to the outcome (Table 1; Supplementary Table S1; Supplementary Figure S11). Assessment for the contribution of covariates to the separation of samples into groups was performed by applying a generalized linear model that employs a maximum likelihood optimization approach (glmnet)^43^. Glmnet combines the advantages of least absolute shrinkage and selection operator (lasso) that tends to select the most correlated covariate and ridge penalty that tends to shrink the coefficients of the correlated covariates, and take advantage of their potentially linear combinatorics. We ran glmnet with lasso and ridge penalty contributing equally to the model (α=0.5) on a 10-fold cross validation mode in order to achieve an optimal set of lambda coefficients. The coefficients proved to be unimportant (Supplementary Table S2; Supplementary Figures S1B, S2B, S9B and S10B). We further investigated whether the detected metabolites were highly associated to covariates by performing Spearman correlation tests while considering the corresponding false discovery rate (FDR) adjusted p-values (Supplementary Table S10).

We first explored the structure of the sets of samples with tissue-specific PCAs (Supplementary Figures S2 and S9). Next we aimed at investigating the separation among phenotypic classes by applying (O)PLS-DA^44,45^. The latter surpasses the requirement of PCA for the variation within the phenotypic groups to be sufficiently smaller than the variation between phenotypic groups. (O)PLS-DA aims at revealing group separations in the score space by employing prior knowledge on the phenotypic group membership while applying an orthogonal signal correction filter to recover for intensity variations that do not correlate to the outcome^42,46^ (Supplementary Figures S1 and S10). (O)PLS-DA tissue-specific model quality was assessed based on the prediction performance Q^2^ statistic, which underperformed due to the limited population (Supplementary Table S3).

### Statistical analysis

We applied non-parametric (Wilcoxon-Mann Whitney test) permutation tests to estimate the statistical importance of the intensities of the metabolites. We used the R package “coin”^47^ that employs Monte Carlo resampling to build the background distribution of the test statistic for a given dataset. We performed 100K permutations and set the threshold for statistical significance to 0.1. We additionally computed the log_2_ fold-change and 90% confidence intervals of the average intensities of metabolites for binary phenotypic groups^7^.

We applied a similar approach to explore the non-monotonic levels of metabolites for the pair-wise phenotypic groups of controls, pre-diabetes and T2D. Specifically, we explored the progress of T2D; hence we focused on the comparison of controls to pre-diabetes, and pre-diabetes to T2D. For the same purpose we also explored linear regression models of HbA1c expressed as a function of various single metabolites and measured their estimation of the proportion of variance explained by the computed regression equation (R^2^) and their p-values. For the linear regression models we used the R package lmPerm that uses permutation tests (100K permutations) for the estimation of statistical significance^48^.

### Multivariate analysis and classification

Additionally, we performed a multivariate variable selection analysis using Boruta^24^. Boruta tackles the problem of selecting *all-relevant* variables that best discern between the phenotypic classes employing random forest classifiers and comparing the relevance of real variables to random synthetic probes. Next we used the selected set of significant metabolites to build tissue-specific classifiers using an R wrapper for the rough-set ROSETTA toolkit^25,49^. ROSETTA computes classifiers in the form of minimal IF-THEN rules that are used to predict an *a priori* defined decision, which in this case was non-diabetes or T2D. In the next step the rules were explored in a network format called VisuNet that facilitates visual exploration of the classifier and demonstrates the co-predictive power of combinations of metabolites^50^.

### Computational analysis tools

To maximize the available information for each metabolite we merged two of the current most widely used small molecules databases, the human metabolome database (HMDB)^51^ and the chemical entities of biological interest (ChEBI)^52^. We also included information from two metabolic-pathway databases, the Kyoto Encyclopedia of Genes and Genomes (KEGG)^53^ and the small molecule pathway database (SMPDB)^54^. Such an integrated database helped us retrieve various information from multiple existing databases for computationally annotated metabolites detected by high-throughput metabolomics technologies (Supplementary Table S11). We implemented a computational tool that converted the information from various metabolomics databases to a unified local collection of data. The tool is available under https://github.com/klevdiamanti/metabolomicsDB.

We additionally implemented a computational tool that performed an integrated analysis of the metabolite intensities, and it is freely available under https://github.com/klevdiamanti/MS_targeted. It takes as input a collection of pre-processed metabolite measurements and their identifiers, and it outputs a set of analysis results that include: i) differential statistical analysis; ii) visual exploration of the statistical analysis; iii) correlation of metabolites and covariates; iv) visual exploration of correlations of metabolites and covariates; vi) datasets for machine-learning approaches; v) correlation analysis for pairs of metabolites; vi) overrepresentation of compounds in metabolic pathways; vii) input datasets for MoDentify^21^; viii) ratio of concentrations for metabolite pairs^55^.

## Data availability

The raw and processed compound levels are available in MetaboLights (https://www.ebi.ac.uk/metabolights/) under the accession number MTBLS690.

## ACKNOWLEDGEMENTS

We would like to acknowledge the Excellence of Diabetes Research in Sweden (www.exodiab.se)in biobank for providing the samples, and the Swedish Metabolomics Centre (www.swedishmetabolomicscentre.se) for the MS and the computational identification of the compounds. The study was supported by grants from AstraZeneca (U.R., J.E., J.K. and C.W.), the eSSence program (J.K.), Institute of Computer Science, Polish Academy of Sciences (J.K.), the Swedish diabetes foundation (C.W.), the family Ernfors fund (C.W.), the Borgström and Hedström fund (M.C.), the Swedish Research Council Formas (M.G.).

## AUTHOR CONTRIBUTIONS

K.D. analyzed the data, interpreted the results and wrote the first draft of the manuscript. M.C. and G.P. prepared the samples, interpreted the results and wrote the manuscript. M.J.P. and J.W.E. interpreted the results and wrote the manuscript. U.R. interpreted the results. M.G. and C.K. assisted in data analysis. S.S. assisted in study design and data analysis. J.K. assisted in the study design, assisted in data analysis, interpreted the results and wrote the manuscript. C.W. conceived the study, assisted in data analysis, interpreted the results and wrote the manuscript. All authors read and approved the manuscript.

